# Life history evolution in response to changes in metapopulation structure in an arthropod herbivore

**DOI:** 10.1101/021683

**Authors:** A De Roissart, N Wybouw, D Renault, T Van Leeuwen, D Bonte

## Abstract

1. The persistence and dynamics of populations largely depends on the way they are configured and integrated into space and the ensuing eco-evolutionary dynamics.
2. We manipulated spatial and temporal variation in patch size in replicated experimental metapopulations of the herbivore mite Tetranychus urticae and followed evolutionary dynamics over approximately 30 generations.
3. A significant divergence in life history traits, physiological endpoints and gene expression was recorded in the spatially and spatiotemporally variable metapopulation, but also a remarkable convergence relative to the stable reference metapopulation in traits related to size and fecundity and in its transcriptional regulation.
4. The observed evolutionary dynamics are tightly linked to demographic changes, more specifically frequent episodes of resource shortage that increased the reproductive performance of mites on tomato, a challenging host plant. This points towards a general, adaptive stress response in stable spatial variable and spatiotemporal variable metapopulations that pre-adapts a herbivore arthropod to novel environmental stressors.

## Introduction

Changes in land-use levy a strong pressure on our natural habitat and leads to habitat loss and isolation, which are both a major thread for biodiversity (Dirzo & Raven 2003). Species conservation therefore relies largely on optimal reserve planning which in turn is rooted within the principles of metapopulations (Levins 1969; Hanski 1998). This concept defines population-extinction dynamics and eventually extinction thresholds within in networks of interconnected habitat patches. Most spatially structured populations can be classified as *patchy* or *mainland-island* metapopulations (Harrison & Taylor 1997), and the omnipresence of classical Levin’s metapopulations has been recently questioned (Fronhofer *et al.* 2012). This spatial variation in habitat availability not only affects patch occupancy dynamics, it also impacts the local- and metapopulation-level demography. Some of us demonstrated experimentally that spatial variation in habitat availability decreases variance in metapopulation size at the metapopulation level (De Roissart, Wang & Bonte 2015). Conversely, spatiotemporal variation in habitat availability increases patch extinction rates, but decreases local population and metapopulation sizes. These demographic changes minimised metapopulation-level variability in mainland-island metapopulations, relative to classical and patchy ones.

Because of these changes in population dynamics, selection pressures in metapopulations are expected to act on more than one level of population structure (Olivieri, Couvet & Gouyon 1990). For instance, in metapopulations where local population extinctions occur regularly, increased dispersal rates are selected relative to metapopulations where extinctions are rare or where patch sizes or heterogeneous, since long-term survival is only possible if genotypes are able to re-colonise patches from where they have become locally extinct. With increasing asymmetry in patch size, however, dispersal will evolve to lower rates because benefits of dispersal are only prevalent for a minority of the individuals (Travis & Dytham 1999). Additional and/or alternative adaptive strategies might also evolve through the adjustment of sex-ratio (Macke *et al.* 2011), age-at-death (Dytham & Travis 2006) and density-dependency (Bierbaum, Mueller & Ayala 1989) according to changes in spatial structure and associated variation in the prevalence and strength of local resource competition (Clark 1978; Cameron *et al.* 2013, 2014) and other stressors (Margulis & Sagan 2000; Parsons 2005).

While evolutionary theory to date is centred on single trait dynamics, multivariate selection in life history and physiology is anticipated in response to changes in spatial habitat configuration. These evolutionary responses then simultaneously feedback on the ecological dynamics, rendering both ecology and evolution heavily intertwined. We now begin to understand such eco-evolutionary dynamics in either natural or experimental metapopulations (Bell & Gonzalez 2011). The importance of eco-evolutionary dynamics is most obvious in metapopulations where dispersal determines the genetic composition and demography of different populations (Kokko & Lopez-Sepulcre 2007). Seminal examples include the Glanville fritillary (Hanski and Mononen 2011) or stick insect metapopulations (Farkas *et al.* 2013). Often, these eco-evolutionary dynamics lead to evolutionary rescue, the process where adaptive evolution allows a population (Gomulkiewicz & Holt 1995), metapopulation (Bell & Gonzalez 2011; Travis *et al.* 2013) or an expanding population (Boeye *et al.* 2013) to recover from negative growth as a result from environmental change (Gomulkiewicz & Holt 1995). Evolutionary rescue is known to be strongly determined by demographic and genetic factors of local populations, but also by entire metapopulation changes (Carlson, Cunningham & Westley 2014).

The genetic basis of life history differentiation can now be disentangled by the development of several –omic approaches. Transcriptomic analyses may uncover genes that significantly alter their transcript levels as a response to the implemented selection pressure and provide detailed insights on the pleiotropic effects underlying phenotypic. For instance, in *Drosophila melanogaster,* many genes that are up- or downregulated in response to stress are equally associated with mobility and aggression (Wheat 2012). In the spider mite *Tetranychus urticae,* transcriptomic analysis of populations that developed pesticide resistance or that were exposed to challenging host plants reveals the presence of common adaptive responses and identified key candidates genes for xenobiotic adaptation in this polyphagous mite (Dermauw *et al.* 2013)

Experimental evolution in artificial metapopulations provides a unique formal test to understand to which degree spatial variation in habitat availability affects life history divergence (Kawecki *et al.* 2012). We installed three types of experimental metapopulation inhabited by spider mites (*T. urticae;* Fig. 1): a stable metapopulation consisting of patches of similar size (the patchy metapopulation; HOM), a metapopulation where habitat quality varies in space and time (the classical metapopulation; TEM) and a metapopulation where patches are of different size (the mainland-island metapopulation; SPA). We earlier documented how this variation in spatial structure affected demographic changes (De Roissart *et al*. 2015) and now report on the evolutionary phenomic changes as measured at the end of that experiment. Following the observed changes in demography and theoretical expectations outlined above, we predicted that:

**Figure 1:**
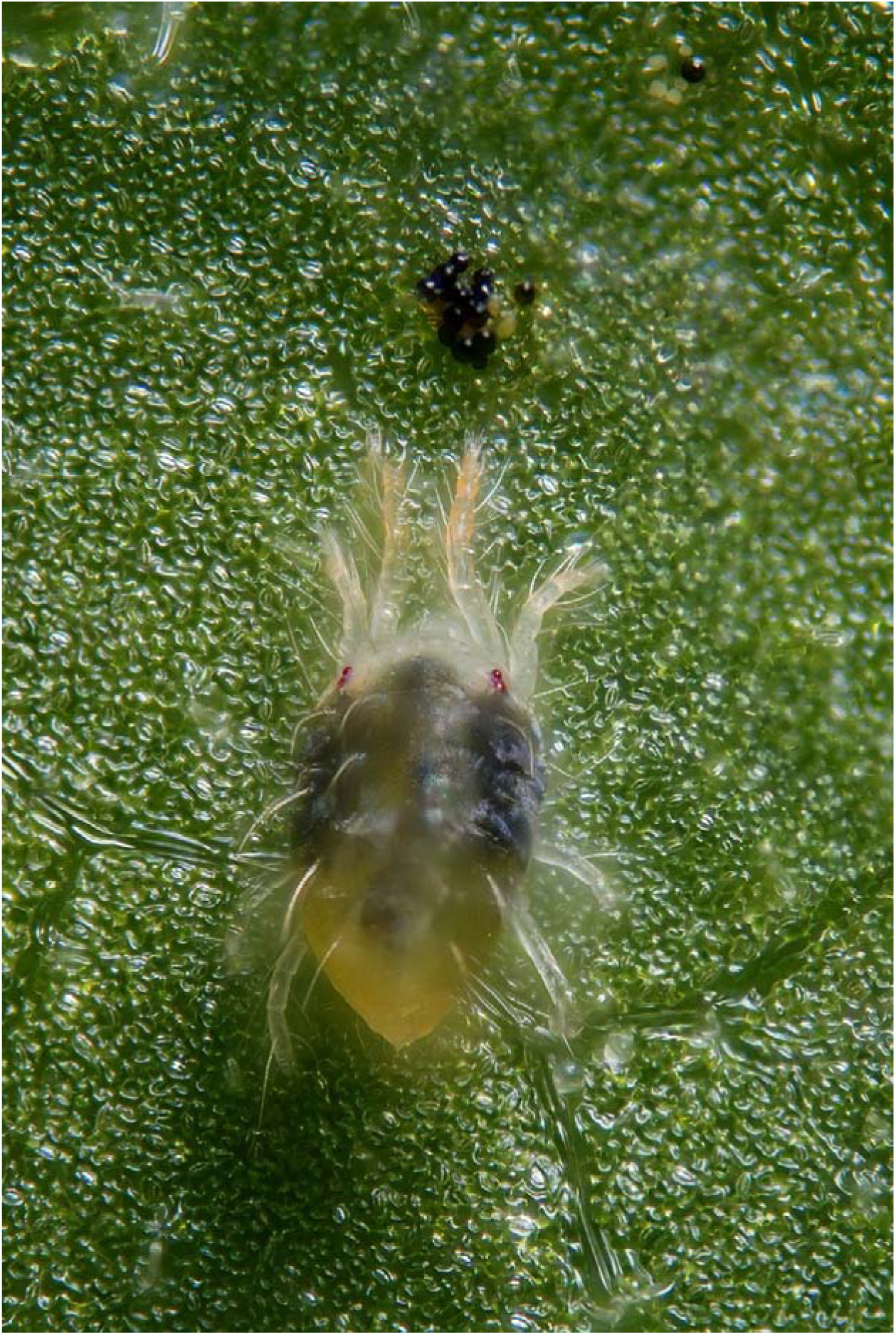
Spider mites as a model for experimental evolution (mature female, on bean). Photo credit by Gilles San Martin

1. Asymmetry in patch and population size in mainland-island metapopulations selects for lower dispersal rates, while in classical metapopulations, the spatiotemporal variation in population size leads to higher evolved dispersal rates
2. Increased local extinction rates and local population variability in classical metapopulation selects for faster life histories, i.e. increased fecundity, decreased longevity and/or reduced developmental time
3. Low local and metapopulation variability in mainland-island metapopulations increase local competition and therefore select for a more male-biased sex ratio and/or increased developmental time (age-at-maturity).

## Materials and methods

### Experimental setup of the artificial metapopulations

Metapopulation dynamics of *Tetranychus urticae* were studied using experimental microcosms. We used as a base population the “LS-VL” *T. urticae* strain, because it is known to be highly evolvable due to its genetic variability (Van Leeuwen *et al.* 2008; Fronhofer, Stelz & Lutz 2014). Artificial metapopulations consisted of a transparent plastic box with 9 patches arranged in a 3 × 3 lattice. We constructed three types of artificial metapopulations with an equal metapopulation-level carrying capacity but varying spatial configuration of the patches. Patches were detached bean (*Phaseolus vulgaris L.*) leaves placed on a Tanglefood layer in closed boxes. This hostile matrix prevents mites from leaving the patches. Bean leaves were renewed on a weekly basis to avoid starvation of the mites. The size of the bean leaves introduced to each patch was dependent upon the treatment. Two times a week, for 8 hours a wind current (1.5m/s) was induced by a fan and allowed aerial dispersal. Three metapopulation types were installed each of which was replicated three times. With exception of the HOM metapopulation, all replicates are similar regarding the specific attributes but different in exact spatial configuration. The three metapopulation types are:

i. a patchy metapopulation consisting of nine patches weekly refreshed with leaves of 20 cm^2^ (spatially homogenous distribution of resources; further referred to as HOM)
ii. a mainland-island metapopulation consisting of three patches of standard leaf size (20 cm^2^) and three of double size; another three patches of these metapopulations remained constantly empty (spatial heterogeneous distribution of resources; further referred to as SPA)
iii. a spatiotemporal heterogeneous metapopulation (further referred to as TEM) in which we assigned nine single-patch resources (standard leaf) randomly to one of the nine patches. Due to this algorithm, the distributions of the resources (and thus local carrying capacity or island size) changed weekly among the nine patches and varied between zero (no resource renewal and local extinction) and double or exceptionally triple island size. In consequence, patch sizes and thus local carrying capacities fluctuated over time and space, but we ensured again a constant metapopulation carrying capacity (9 × 20 cm^2^) over time.

At the beginning of the experiment, 20 randomly collected adult female mites, from the base population, were assigned to each patch within each metapopulation type and allowed to establish the triplicated populations. All metapopulations were kept under controlled conditions (23°C, 16:8 LD photoperiod, 85% humidity).

### Quantification of mite life-history

Spider mite life-history traits were measured at the initiation of the experiment and after 10 months, corresponding to approximately 30 mite generations. All traits were measured on F2 mites (raised for two generations in common garden on detached leaf discs) to minimise maternal and environmental effects caused for instance by local conditions of crowding. Young inseminated females of each experimental metapopulation were individually allowed to oviposit on bean leaf discs. Leaf discs were placed with the abaxial part upwards on moistened filter paper to prevent mites from escaping and to maintain leaf turgor. Different life history parameters of the descendants were recorded daily: juvenile survival, developmental time (time from egg until the adult stage), fecundity (daily number of eggs), adult longevity and sex-ratio. Since spider mites deposit the majority of their eggs during the first seven days after maturity, we monitored fecundity only during that period. Dispersal propensity of the mites was assessed by transferring mated females to test arenas for trials of aerial dispersal (after two whole generations under common garden to avoid confounding maternal effects). The experimental setup for aerial dispersal assessment was identical to the one applied in (Li & Margolies 1994) (details in Appendix S1 in supporting information).

### Mite performance

Mite performance was followed by quantifying rate of intrinsic growth as a proxy of fitness (Cameron *et al.* 2013). To detect possible differences in individual performance between treatments, an integrated individual-level fitness measure, the rate of intrinsic growth (r_m_), was calculated by combining the estimated parameter distributions of the different life history parameters according (see statistical analyses) to the equation 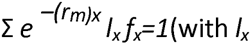 (with *I_x_* survival till maturity *x* and *f_x_* the number of female offspring at age *x*) which represents the contribution of each female to the number of females in the subsequent generation. We performed 10000 simulations and reported the mean value and standard deviation while testing its significance in comparing whether 2.5% tails of the distribution overlap. We additionally measured a set of physiological endpoints (mass, glucose, trehalose and triglycerid levels) after the common garden treatment at the start of the experiment and after 30 generations of selection in a metapopulation context as indicators for mite performance. All physiological parameters were measured following (Laparie *et al.* 2012) on F2 mites (Appendix S1 in supporting material).

### Differential gene expression after experimental evolution

To examine the effects of metapopulation structure on the mite transcriptome, Agilent dual colour gene expression micro-array analysis was performed on female mites raised for two generations in a common garden of every selection regime. The microarray data have been deposited in the Gene Expression Omnibus (GEO) (accession number: GSE55623). For the hierarchical clustering, data of previous *T. urticae* studies were incorporated (Bryon *et al.* 2013; Zhurov *et al.* 2014). Final statistical processing and analysis was conducted in limma (Smyth 2005). Gene Ontology (GO) annotation was executed using Blast2GO software (Conesa *et al.* 2005). Using the Blast2GO generated annotation and the statistical output of limma as input, Gene Set Analysis (GSA) was performed with the Bioconductor package piano (Parametric Analysis of Gene set Enrichment, PAGE)(Väremo, Nielsen & Nookaew 2013). More details of the gene expression and GO-term analysis are provided in Appendix 1 from Supporting Material.

### Performance on a challenging new host

Our LS-VL base population has been maintained on bean for more than 10 years. We assessed performance on a novel suboptimal host by quantifying isofemale growth rate on tomato (*Solanum lycopersicum;* variety Moneymaker) grown under controlled laboratory conditions (23°C, 16:8 L:D photoperiod). Experimental arenas were constructed with leaves from 4-week old tomato plants. Moist tissue paper was used to cover 10 cm^2^ leaf edges that prevented mites from escaping. Twenty fertilized F2 females (raised for two generations in common garden to reduce maternal and environmental effects) from each artificial metapopulation were placed on a leaf-arena and allowed to establish a population. All leaf-arenas were kept under controlled conditions (23°C, 16:8 L:D photoperiod). Population growth was assessed weekly for 3 weeks by counting the number of eggs, juveniles, adult males and females.

### Statistical analysis

We first tested for overall multivariate divergence in life history after experimental evolution and subsequently used GLMM to test the univariately according to the imposed treatments. The measured traits follow different statistical distributions and therefore applied a Permutational Multivariate Analysis of Variance (PERMANOVA). Because our measurements were taken with different units on different scales, the correctly estimated replicate-level averages of the life history and physiological endpoints (see GLMM further) were scaled prior to PERMANOVA analysis based on Euclidean distances among replicates belonging to one of the three metapopulation treatments (PERMANOVA; with adonis function in R; (Anderson 2005). To visualise metapopulation divergence based on life history, Nonmetric Multidimensional Scaling (NMDS) analyses were performed on the scaled distance matrix (all life history and physiological traits) using the metaMDS function (vegan library, R.2.15.1;). The significantly diverging traits were subsequently identified by a Multivariate Analysis of Variance (MANOVA) on the scaled averaged data per replicate.

We examined how metapopulation type affected the different life history traits and physiological endpoints using generalized linear mixed models (GLMM). The model included metapopulation type (HOM, SPA, TEM) as fixed factor and each individual metapopulation as a random effect to control for dependency among the three replicates from each metapopulation treatment. Depending on the dependent variable, a Gaussian (all physiological endpoints), Poisson (fecundity, developmental time, longevity and population size on the novel host) or binomial error (sex ratio, juvenile mortality) structure was modelled with appropriate link functions. Non-significant contributions (P>0.05) were removed by backwards procedure. Effective degrees of freedom were estimated using Kenward-Rogers procedure. All analyses were conducted with SAS 9.3 (SAS Institute Inc 2006) by using the GLIMMIX procedure.

## Results

### Population-level divergence in life history traits

Experimental evolution caused significant divergence in life history traits between the treatments (permanova F_2_= 2.75; p=0.03). MANOVA analyses showed sex-ratio (F_2_=7.77; p=0.02) and fecundity (F_2_=10.35; p=0.01) as the two main life history endpoints underlying this divergence. A detailed analysis on the individual trait distribution after experimental evolution confirmed divergence in fecundity and sex ratio, but also in longevity (Table 1; Fig 2). An analysis of the trait variation at the start of the experiment is provided in Appendix S2 from the supporting material.

**Table 1:** Results for fixed effects from mixed linear models with fecundity, developmental time, sex-ratio, longevity and juvenile survival as response variable.

| Factor | Num df | Den df | *F* | *p* |
| --- | --- | --- | --- | --- |
| ***Daily fecundity*** |  |  |  |  |
| Treatment | 2 | 8.732 | 9.34 | 0.0068 |
| Day | 1 | 1891 | 693.77 | <.0001 |
| Day*Treatment | 2 | 1891 | 2.99 | 0.0507 |
| ***Total fecundity*** |  |  |  |  |
| Treatment | 2 | 106.2 | 6.79 | 0.0017 |
| ***Sex-Ratio*** |  |  |  |  |
| Treatment | 2 | 189.8 | 3.55 | 0.0308 |
| ***Developmental time*** |  |  |  |  |
| Treatment | 2 | 2228 | 4.09 | 0.0168 |
| Sex | 1 | 228 | 10.78 | 0.0010 |
| Treatment*Sex | 2 | 2228 | 0.25 | 0.7782 |
| ***Longevity*** |  |  |  |  |
| Treatment | 2 | 158 | 5.43 | 0.0053 |
| ***Juvenile survival*** |  |  |  |  |
| Treatment | 2 | 7.201 | 0.25 | 0.7882 |

**Figure 2:**
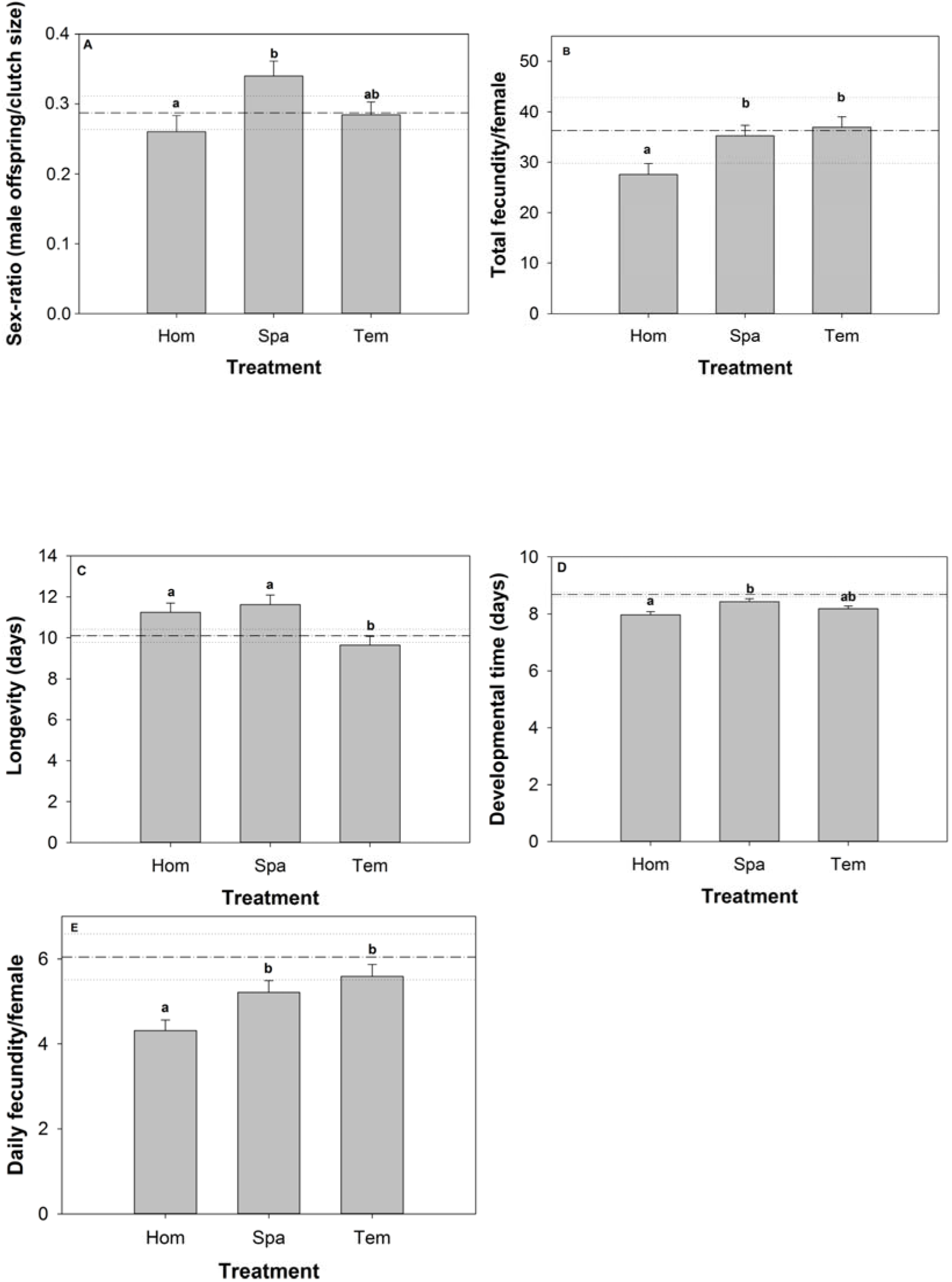
Effects of variation in metapopulation structure on life history parameters (mean values ± SE) of mites. A: longevity, B: total fecundity, C: sex ratio (males/total clutch size), D: developmental time; E: Daily fecundity. Dotted lines represent parameter values before 30 generations of selection. Equal notations indicate non-significant contrast for the respective measurements. Error bars represent standard errors.

The average proportion of male offspring was higher in clutches originating from the SPA metapopulations (0.34 ± 0.02SE) relative to the HOM (0.26 ± 0.02SE). Mites from the SPA and TEM treatment evolved a high daily fecundity (t=−3.79; p=0.01) than those from the HOM treatment (respectively 5.21± 0.28SE and 5.59± 0.28SE versus 4.31± 0.25SE). Similarly, mites that evolved in the SPA (35.20± 2.01SE) and TEM treatment (36.91± 2.1SE) had a significantly higher fecundity (t=−3.53, p=0.0014) than those from the reference population HOM (27.61± 2.01SE).

Mites that evolved in the TEM metapopulations died earlier after reaching maturity (9.65 ± 0.42SE days) than mites from homogeneous (HOM) metapopulations (after 11.24 ± 0.45SE days) and spatial variable (SPA) metapopulations (after 11.62 ± 0.47SE days) (Fig. 2C). Under the prevailing lab conditions, males developed in 7.99 days on average while the female reached maturity after 8.40 days (t=−3.28; p=0.0010). Mites from homogeneous metapopulations reached maturity earlier (7.97 ± 0.12) than mites from the spatially heterogeneous (SPA) metapopulation (8.43 ± 0.11) (t=−2.85; p=0.012) (Fig. 2D). The interaction between sex and treatment was not significant (F_2_,_2228_=0.25; p=0.78). No significant differences in juvenile survival of mites among treatments were observed (F_2_,_7.201_=0.25; p=0.79), and no differences were detected in aerial dispersal propensity (F_2,5.185_=0.02; p=0.98).

The simulated growth rate at the start of the experiment was 3.56 (SD=0.19). After experimental evolution, growth rates were slightly lower in the homogeneous metapopulation treatment relative to the other two, but this difference was not significant based on the inferred 95% confidence intervals (r_Hom_=3.38, SD=0.21; r_TEMP_=3.52; SD=0.20; r_SPA_=3.55, SD=0.19).

### Divergence in physiological endpoints

Although not significant (F_2_,_32_= 3.08; p=0.06), a trend towards a lower mass per 50 mites was observed for mites from homogeneous metapopulations (424 ± 25SE μg) compared to mites from metapopulations with spatial (510 ± 25SE μg) or spatiotemporal variation (441 ± 31SE μg). Glucose levels were significantly different among the metapopulation treatments (F_2_,_67_=3.52; p=0.03; Fig. 3), with the lowest levels for HOM (1.39 ± 0.25SE) relative to those from SPA (2.33 ± 0.25SE) (t=−2.64; p=0.027). No significant differences in trehalose (F_2_,_60_=0.43; p=0.51) or triglyceride level were observed among treatments (F_2_,_56_=2.07; p=0.14).

**Figure 3:**
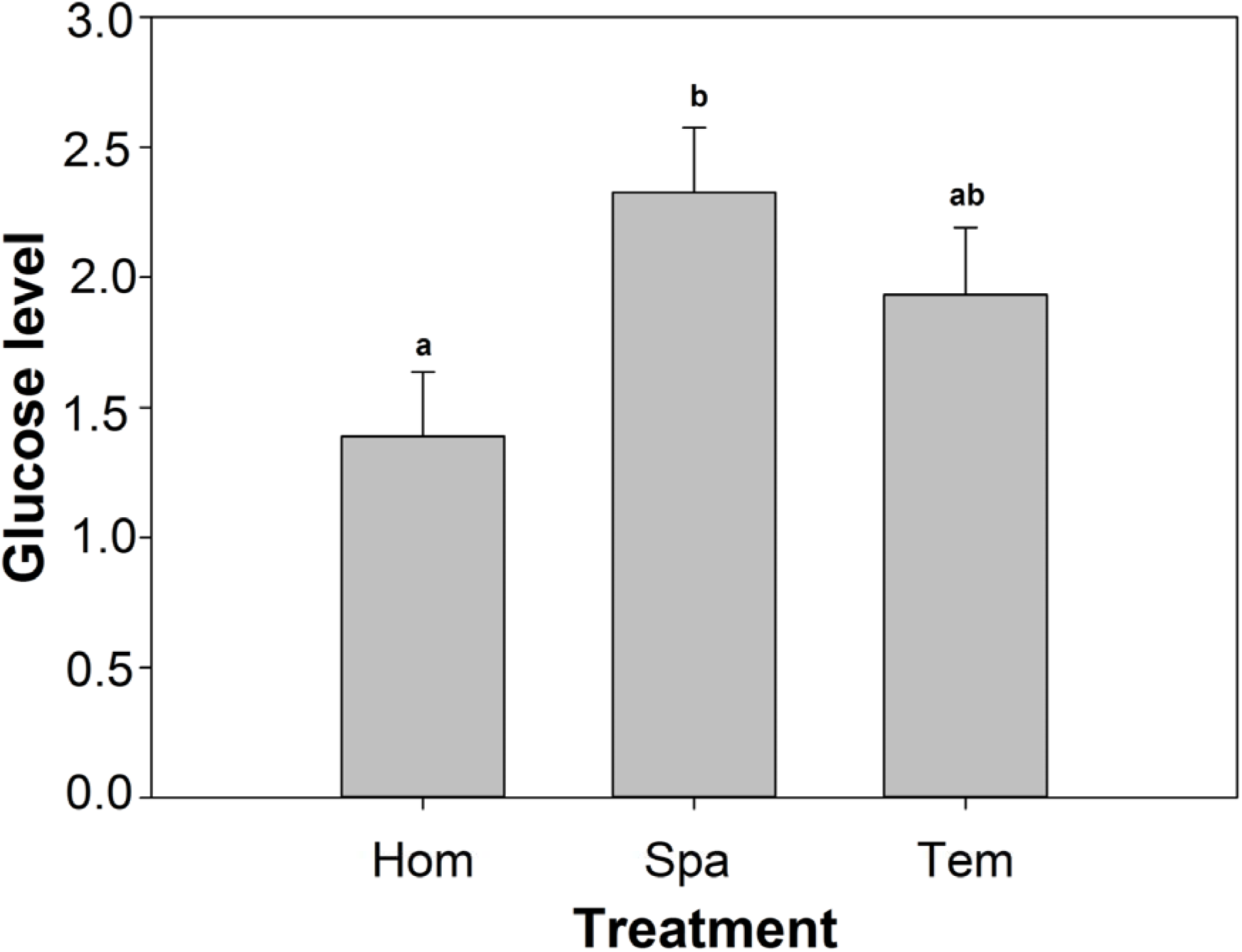
Effects of variation in metapopulation structure on glucose level (nmol) per 50 mites (mean values ± SE). Equal notations indicate non-significant contrast for the respective measurements. Error bars represent standard errors.

### Divergence in gene expression

Based on genome-wide gene-expression data of adult female mites raised under common garden for two generations, SPA and TEM treatments diverged from the control HOM regime, in a parallel direction. We found 152 and 181 differentially expressed genes in SPA and TEM lines, respectively, using the HOM treatment as reference (FDR-corrected *p*-value <0.05 and log_2_-converted fold change (FC)>0.585) (Fig. 4). Fig. S1 depicts the expression patterns of the triplicated lines within each treatment separately. Of these differentially expressed genes, 81.6% and 70.7% exhibited down-regulation in SPA and TEM, relative to HOM, respectively (Fig. 4, Fig. S1 in supporting material). Pearson correlation indicated that the altered transcript levels in SPA and TEM were significantly correlated (ρ=0.80, df=260, p<0.0001).

**Figure 4:**
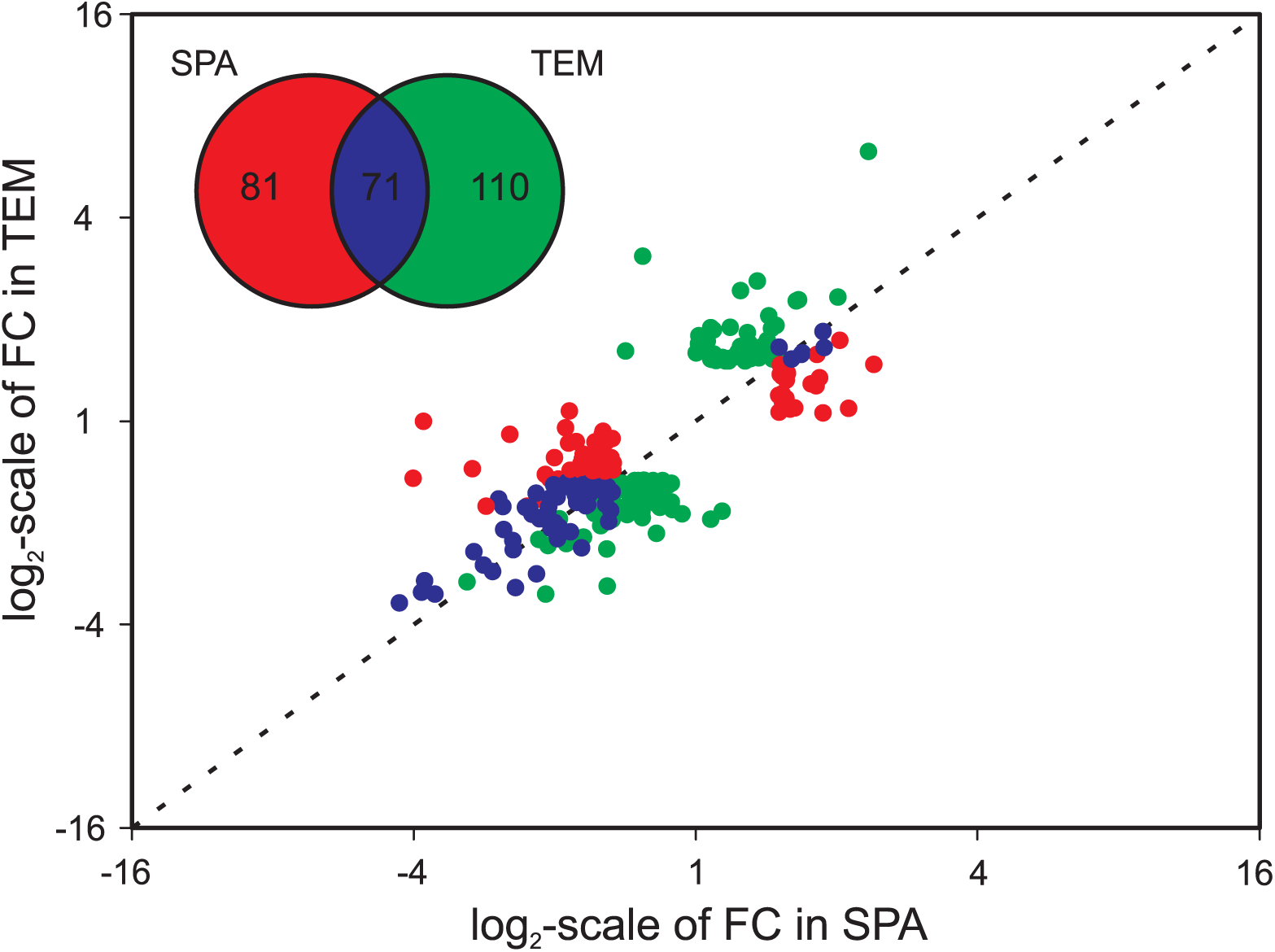
Scatterplot showing the log_2_-scaled fold change of the differentially expressed genes in SPA and TEM, using HOM as reference. The inset represents a Venn-diagram depicting the number of differentially expressed genes in the TEM and SPA lines, relative to HOM. The labels on the log_2_-scaled x- and y-axis represent the non-transformed fold change (FC) values.

Using Blast2GO (Conesa *et al.* 2005), a total of 164 Biological Process GO-sets were assigned to the differentially expressed genes that were associated with evolution in the TEM and SPA regimes. Approximately half of these GO-terms (n=84) were present in both treatments. Twenty gene sets were significantly up- and down-regulated in either the SPA or TEM regimes, relative to HOM (Fig. 5). The associated labels of the GO-IDs are listed in Table S1 in supporting Material. Using this GSA, we observed a convergence in the down-regulation of genes involved in methionine biosynthesis (Fig. S3 in supporting material). In contrast, the transcriptional responses to TEM and SPA selection pressures diverged in the gene sets of which members were associated with gluconeogenesis and interconnected pathways and in gene sets of which members code for glycoside hydrolases (Fig. 5 and Fig. S2 in supporting material).

**Figure 5:**
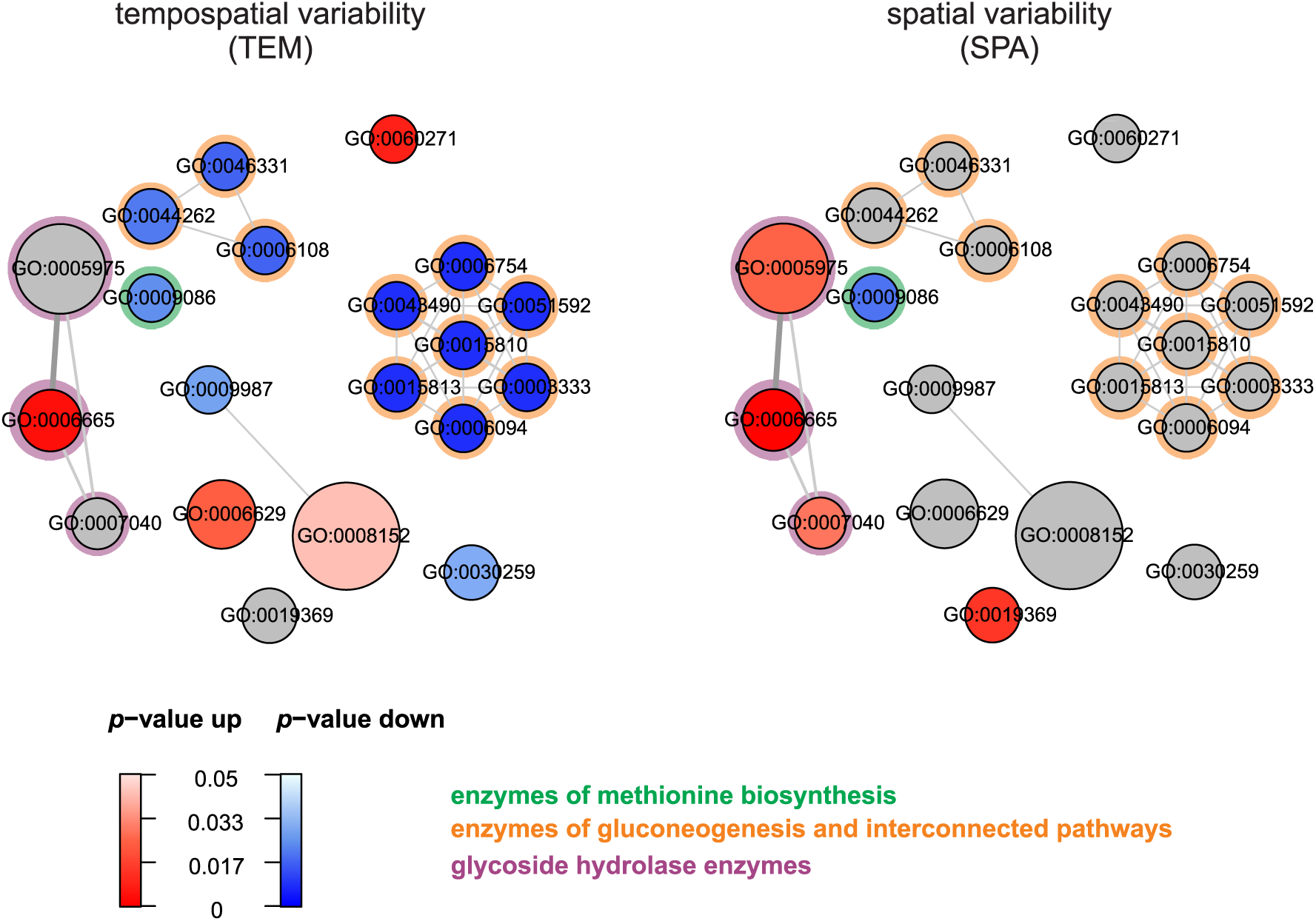
Gene set analysis of biological processes for differentially expressed genes in spider mites after adaptation to spatial and spatiotemporal variability. Nodes and edges represent gene sets and overlap of members between interconnected sets, respectively. Using PAGE, red, blue and grey indicate whether a gene set was significantly up-regulated, down-regulated or not differentially expressed, respectively (Varemo et al., 2013). Gene sets are labelled with the GO-ID (corresponding GO-labels are listed in Table S1 in the supporting material). Green, orange and purple halos surrounding nodes show which gene sets code for genes involved in methionine biosynthesis, gluconeogenesis and interconnected pathways, and genes coding for glycoside hydrolases, respectively.

### Population performance on novel hosts

After one week of challenging the novel host, mite survival did not differ according to the spatial setting to which they evolved (F_2_,_122_=2.19; p=0.12). However, significant differences in fecundity were observed (F_2_,_122_=66.81; p<0.0001), with a lower number of deposited eggs in mites that evolved in the homogeneous populations (49.33 ± 1.07SE eggs) relative to SPA (68.29 ± 1.29SE eggs) and TEM (55.83 ± 1.17 eggs). All pairwise differences were significant (Fig. 6A). After three weeks, the first cohort of offspring matured which differed in population size among treatments (F_2_,_5.635_=5.83; p=0.04; Fig. 6B). Again, population sizes were lowest in mites originating from HOM (5± 0.90SE) relative to SPA (10.26 ± 1.71SE) and TEM (10.62 ± 1.77SE).

**Figure 6:**
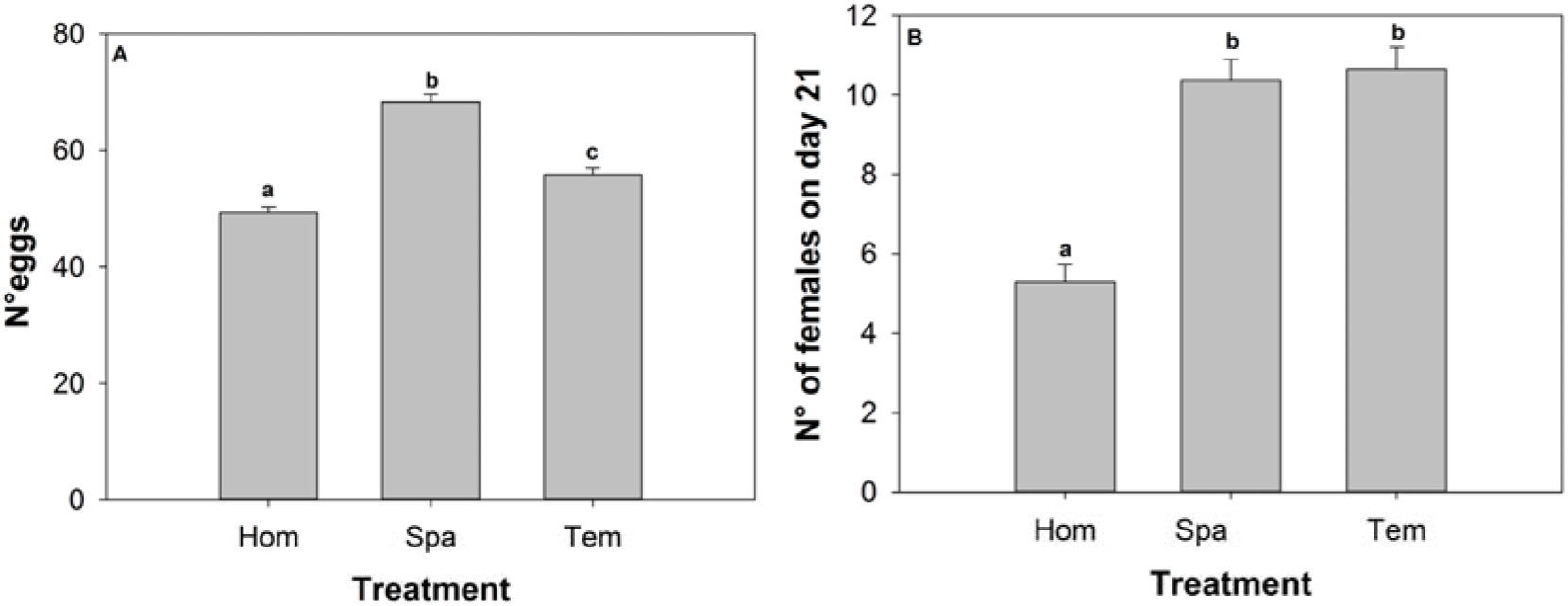
Effects of long-term evolution in the different metapopulation contexts on population growth on a novel host (mean values ± SE). A: number of eggs after one week, B: number of female offspring reaching adulthood after 21 days. Equal notations indicate non-significant contrast for the respective measurements. Error bars represent standard errors.

## Discussion

While there is an increasing awareness that changes in spatial structure affect population dynamics, and that these ecological dynamics interact with evolutionary trajectories, there is a limited understanding of how these reciprocal eco-evolutionary interactions are governed by metapopulation-level selection pressures. We followed a phenomic approach in which we contrasted life history traits, physiological endpoints and transcriptomes from mites that evolved in classical and mainland-island metapopulations with those that evolved in stable patchy metapopulations. Using stable patchy metapopulations (HOM) as a reference, mites from the spatially (mainland-island, SPA) and spatiotemporally (classical, TEM) variable metapopulations showed evolutionary convergence in traits related to size and fecundity and in its transcriptional regulation of methionine biosynthesis. However, these mites equally evolutionary diverged in longevity, sex ratio, glucose content and in the expression of genes involved in gluconeogenesis and that code for glycoside hydrolases (Fig. S2 in supporting information). Despite this divergence in the separate life traits, overall population growth rate on the ancestral host did not differ among the treatments because of different population-level trade-offs.

Relative to HOM, spatiotemporal variation in habitat availability in the classical metapopulations (TEM) generated high levels of local variation in population density and on average lower metapopulation sizes due to frequent patch extinctions and lagged colonisation dynamics (De Roissart *et al*. 2015). Mites evolving in these metapopulations showed a significant down-regulation of transcription of genes associated with gluconeogenesis and ATP production (Fig. S2). Genes of the gluconeogenesis pathway affect various metabolic fluxes and energy production and are known to influence life history traits (e.g. dispersal, life span and basal metabolic rate), which determine survival in a metapopulation structure. Here, mite adaptation to spatiotemporal dynamics led to an increased fecundity and reduced adult longevity, which may have been mediated by the transcriptional changes in their gluconeogenesis pathway (Fig S2 in supporting information). These altered traits and resulting trade-off reflect a change in resource allocation between survival and reproduction (Magalhães *et al*. 2007), leading to the evolution of more *r-*strategic traits (Ronce, Perret & Olivieri 2000; Wheat & Hill 2014). In general, the evidence for the existence of such trade-offs is poorly documented.

Mainland-island metapopulations (SPA) are characterised by both a low local and metapopulation-level variability in population size, large metapopulation sizes and low metapopulation-level dispersal (De Roissart *et al*. 2015). We detected no such pattern at the individual level. This contrasts with theoretical predictions (Ronce *et al*. 2000) and can be explained by the densities in the trials at which differences are not expressed, the behavioural lability which hides potential minor average differnces, or the overruling impact of kin competition in our system that hides potential minor differences in this behaviour. More importantly, however, an evolution towards more male biased, fecund and slowed aging strategies relative to mites from patchy metapopulations (HOM) was detected. Sex ratio changes are known to evolve in response to local resource (Clark 1978) or mate competition (Macke *et al.* 2011). While mites from stable metapopulations with low dispersal and stable local population sizes are expected to evolve more female biased sex-ratios due to elevated kin-competition, our results point in the direction of resource competition. Spider mites show scramble competition, with fast resource depletion when population sizes are high (Krips *et al.* 1998). As such, slowing down population growth by extending age-at-maturity (Cameron *et al.* 2014; Monro & Marshall 2014) and more male-biased sex ratios (Johnson 1988; Zhurov *et al.* 2014) can be considered as an adaptive strategy under stable conditions and elevated resource competition. The significantly higher transcription in gene sets that code for glycoside hydrolases, enzymes which are crucial for the digestion of complex carbohydrates in an arthropod herbivore’s diet and consequently a higher feeding efficiency (Terra & Ferreira 1994), can be regarded as a consequence of mite adaptation to the stable conditions and the resulting elevated resource competition. Interestingly, two of the differentially expressed *GH* genes (*tetur29g01280* and *tetur29g01230*) code for glycoside hydrolases of the GH32 family and seem to have been incorporated in the mite’s genome by a horizontal gene transfer event from a bacterial donor species (Grbić *et al*. 2011). The mite genome harbours a remarkable number of horizontally transferred genes and the observed evolutionary response in the transcription of these *GH* genes to metapopulation selection seems to support the hypothesis that horizontal gene transfer is a driving force in the adaptive evolution of spider mites (Grbic et al. 2011).

Despite the evolved divergent phenotypic profiles reported above, life history traits and genome-wide gene-expression showed an overall convergence in the spatial and spatiotemporal variable metapopulation configurations, relative to the stable patchy one. The convergence is apparent in (a) similar evolution of higher fecundity rates and to a lesser degree sex-ratio, age and mass-at-maturity, (b) and increased glucose content and (c) the identical direction of the differential expression of 71 genes (including genes of the methionine anabolic pathway; see Fig. S3 in supporting information). Increased fecundity shows trade-offs with longevity and sex ratio in respectively the TEM and SPA metapopulation. Therefore, mites did not evolve an increased per capita growth rate and on average reached maturity at later age. A slower growth, is typically associated with competitive environments (Kawecki 1993). For the SPA treatment, such increased competitive interactions have been discussed above, but densities were on average lower in the TEM treatments (De Roissart *et al*. 2015), rendering high densities as a common competitive environment a poor explanation for the detected convergence. Systematic high densities in the SPA treatments and resource limitation combined with unoccupied patches in the TEM treatment (De Roissart et al. 2015) must have led to more frequent episodes of *per capita* resource shortage, relative to the HOM metapopulations. Elevated glucose levels have been associated with responses to cope with increased starvation resistance in a ground beetle (Laparie *et al.* 2012) and are typically associated with low metabolic rates under food limitation(Packard & Boardman 1999; Božič & Woodring 2015). A slower development and higher glucose levels of mites from the SPA and TEM are thus in concordance with evolutionary trajectories towards stress resistance (Sulmon *et al.* 2015).

Our theory of a general stress response was further supported by the transcriptional responses. First, identical expression profiles of a GO-term related to methionine synthesis indicates a common response that could interfere with methylation processes. Such methylation is often induced by general, oxidative stresses, for instance –and in line with our findings-due to compensatory growth after food restriction (De Block & Stoks 2008). Moreover, the more frequent resource shortage experienced by TEM and SPA populations may have restricted their access to sulfur-rich proteins containing methionine from *P. vulgaris* (George *et al.* 1993). During the common garden experiment, the increased access to the beans for SPA and TEM may have reduced the need for methionine synthesis, an amino acid that has been proven a key player in driving fecundity in *Drosophila melanogaster* (Zajitschek *et al.* 2013).

Significantly up-and down-regulated gene sets upon adaptation to (spatio)temporal stress included sets that are associated with basal metabolic pathways. Genetic changes in these pathways are a common response to environmental stressors, with enzymes of the gluconeogenesis/glycolysis and citric acid pathways as one of the prime targets (Marden 2013). It is interesting to notice that glucose 6-phosphatase shares a substrate with PGI (see Figure S2 in supporting material), a protein previously connected to performance in specific metapopulation structures in butterfly species (Wheat & Hill 2014). It may therefore represent an important link between stress resistance and dynamics in metapopulations.

Adaptive metabolic changes are known to lead to the development of cross-tolerance in organisms, enabling organisms to cope with unfamiliar stressors. We demonstrated that evolutionary dynamics resulting from changes of the metapopulation spatial structure, pre-adapt mites to cope with a challenging novel host. This is an important finding which definitively needs more study in other (model) organisms. If general, such an evolutionary response is expected to have a strong impact on community- and food web dynamics under natural conditions (Farkas *et al.* 2013). We show that altered population dynamics due to changes in metapopulation spatial structure may induce general stress resistance responses. Since multiple stressors are jointly operational under global change, evolutionary responses towards changes in spatial structure and the coupled spatiotemporal variation in demography may offset the need for adaptation to other environmental stressors by maintaining a general stress response and lead to evolutionary rescue (Carlson *et al*. 2014).

## Acknowledgements

This project was funded by FWO projects G.0610.11 and G.0093.12N and Belspo-IAP project Speedy. ADR was funded by BOF-Ugent. NW was supported by the Institute for the Promotion of Innovation by Science and Technology in Flanders (IWT, grant IWT/SB/101451). DR was supported by the Observatoire des Sciences de I’Univers de Rennes (OSUR).

## Author contributions

ADR, DB & TVL designed the study, ADR & NW performed the research, ADR wrote the first draft of the manuscript and analyzed the data, and all authors contributed substantially to revisions.

## Supporting information

Additional supporting information may be found in the online version of this article.

Appendix S1: additional information methodologies

Appendix S2: trait variation at the start of the experiment

Table S1: GO labels

Fig S1: expression heatmap

Fig S2: citric acid cycle, glycolysis and gluconeogenesis

Fig S3: methionine synthesis pathway

